# A composite frailty index enables quantification of functional aging and identification of gerotherapeutic drugs in the house cricket

**DOI:** 10.64898/2026.04.01.715973

**Authors:** M.S. Gerald Yu Liao, D.V.M. Jenna Klug, B.S. Swastik Singh, D.V.M. Warren Ladiges

## Abstract

Frailty, defined by progressive loss of physiological resilience, neuromuscular function, and cognitive capacity, is a central manifestation of biological aging yet remains difficult to quantify in scalable experimental systems. Here, we introduce a Composite Frailty Index (CFI) in the house cricket (*Acheta domesticus*) that integrates automated measures of locomotion, exploratory behavior, and freezing into a unified, quantitative framework of functional decline. Ten behavioral parameters derived from automated open-field tracking, including locomotor performance, exploratory behavior, and freezing were integrated into the CFI. Locomotor states were classified using k-means clustering (*k* = 2) of velocity distributions, and all features were normalized to age- or treatment-matched reference populations, discretized into quintiles, and summed to generate a 0-40 frailty score. Aging cohorts (young adult: 4-6 weeks; geriatric: 10-12 weeks, N = 103) and pharmacological cohorts treated at mid-life (8-10 weeks) with rapamycin (14 ppm), acarbose (1000 ppm), or phenylbutyrate (1000 ppm) were evaluated (N = 122). Across chronological aging cohorts, CFI increased from young adults to geriatrics in both females (*d* = 1.14 [95% CI: 0.53, 1.76], *P* = 0.0003) and males (*d* = -1.17 [95% CI: -1.75 to -0.59], *P* < 0.0001). Using pharmacological intervention cohorts, mid-life rapamycin treatment reduced late-life frailty relative to controls in both females (*d* = -1.31 [95% CI: -2.09, -0.53], *P* = 0.0017) and males (*d* = -1.33 [95% CI: -2.09, -0.58], *P* = 0.0004), whereas acarbose and phenylbutyrate produced inconclusive effects (*d*’s = -0.54 to -0.03; *P*’s > 0.05). Together, these findings establish the cricket CFI as a scalable, high-throughput platform for quantifying multidimensional functional aging and prioritizing candidate geroprotective interventions based on clinically relevant endpoints beyond lifespan.

## Introduction

Frailty is a conserved biological state characterized by progressive loss of physiological resilience, impaired neuromuscular function, and reduced cognitive capacity, leading to increased vulnerability to disease and mortality [1]. In humans, frailty predicts adverse outcomes across organ systems independent of chronological age, positioning it as a central phenotype of biological aging rather than a secondary clinical condition [2]. Accordingly, preserving functional capacity has emerged as a primary goal of geroscience, as interventions that delay frailty may reduce the burden of multiple age-related diseases simultaneously [3].

Despite this shift, most preclinical studies still rely on lifespan extension as the principal endpoint [4]. While survival-based screens have identified candidate geroprotective compounds, they provide limited insight into functional decline, the defining feature of frailty and a key determinant of quality of life [5]. Frailty reflects coordinated deterioration across physical performance, neuromuscular integration, and behavior, which can diverge from survival outcomes, highlighting a critical gap between experimental metrics and clinical reality. Scalable platforms capable of quantifying multidimensional functional aging are therefore needed [6].

Invertebrate models offer high throughput and experimental control but remain underutilized for frailty research due to perceived limitations in physiological and behavioral complexity [7-8]. The house cricket (*Acheta domesticus*) provides a complementary system with defined neuromuscular and neural circuits and measurable age-dependent declines in locomotor performance, coordination, and behavioral adaptability [9-12]. Here, we develop a Composite Frailty Index (CFI) integrating automated measures of physical and cognitive function into a unified quantitative framework for high-throughput assessment of functional aging. Using this platform, we evaluate FDA-approved geroprotective drugs (rapamycin, acarbose, and phenylbutyrate) which offer translational potential due to established safety profiles and mechanisms [13], testing their ability to attenuate age-associated increases in frailty and preserve functional capacity.

## Methods

### Animals

House crickets (*Acheta domesticus*) of heterogeneous genetic background were obtained from a commercial supplier (Fluker Farms Inc, Louisiana, USA) and maintained under standardized conditions (29 ± 1°C and 32 ± 3% relative humidity) optimized for physiological stability [9,11]. Animals were group-housed (20-30/cage; mixed sex) in a double-enclosure system permitting self-regulated photoperiod exposure, with egg carton shelters to facilitate light avoidance and naturalistic behavior [11,14]. Crickets were not maintained under specific pathogen-free conditions to preserve naturalistic aging trajectories [15]. All animals underwent ≥1 week acclimation prior to experimentation [16].

For aging analyses, crickets were stratified by post-eclosion age into young adults (4-6 weeks) and geriatrics (10-12 weeks) (N_Young Adult_ = 30 of each sex, N_Geriatric_ = 19 female, 24 male), corresponding to preserved versus impaired functional states [11]. For pharmacological studies, cohorts were randomized at 8 weeks (mid-life, corresponds to when moderate functional decline begins [17]) to control (N = 28 female, 34 male) or treatment groups (N = 10/sex/group, total = 60), treated daily for two weeks, and assessed at 10 weeks.

### Diet preparation and drug administration

Crickets were maintained on a standardized diet of irradiated rodent chow (Picolab Rodent Diet 20, 5053; Purina Mills, USA) incorporated into a gelatin matrix to ensure microbial stability and uniform drug delivery. The diet was prepared by dissolving gelatin, mixing with powdered chow, cooling, dehydrating, and homogenizing [18]. Water was provided via gel packs (Napa Nectar Plus) and replaced every 48 h.

For chronological aging cohorts, food was provided *ad libitum*. For pharmacological cohorts, intake was controlled (2 g/cage every 72 h) with regular replacement to maintain compound stability. Drug-supplemented diets were formulated using the same base matrix at dosages established in murine studies [19], with preparation adjusted for compound-specific stability and solubility [18]. Rapamycin (14 ppm; Southwest Research Institute, San Antonio, TX) was directly incorporated into the gelatin-chow matrix, whereas phenylbutyrate (1000 ppm; Triple Crown America Inc., Perkasie, PA) and acarbose (1000 ppm; Spectrum Chemical Mfg Corp, Gardena, CA) were pre-dissolved in water prior to incorporation. Control groups received identical un-supplemented diet.

### Open field behavioral assay and automated tracking

Locomotor and exploratory behaviors were assessed using a 5-min open field assay conducted in darkness (Fig. 1A). Crickets were placed at the arena center, video-recorded, and arenas were sanitized with 10% bleach between trials. Recordings were analyzed using ezTrack [20-21], with positional coordinates and velocity extracted at 30 fps following arena calibration. The arena was partitioned into central and peripheral zones to quantify spatial behavior (Fig. 1B).

**Fig. 1.**
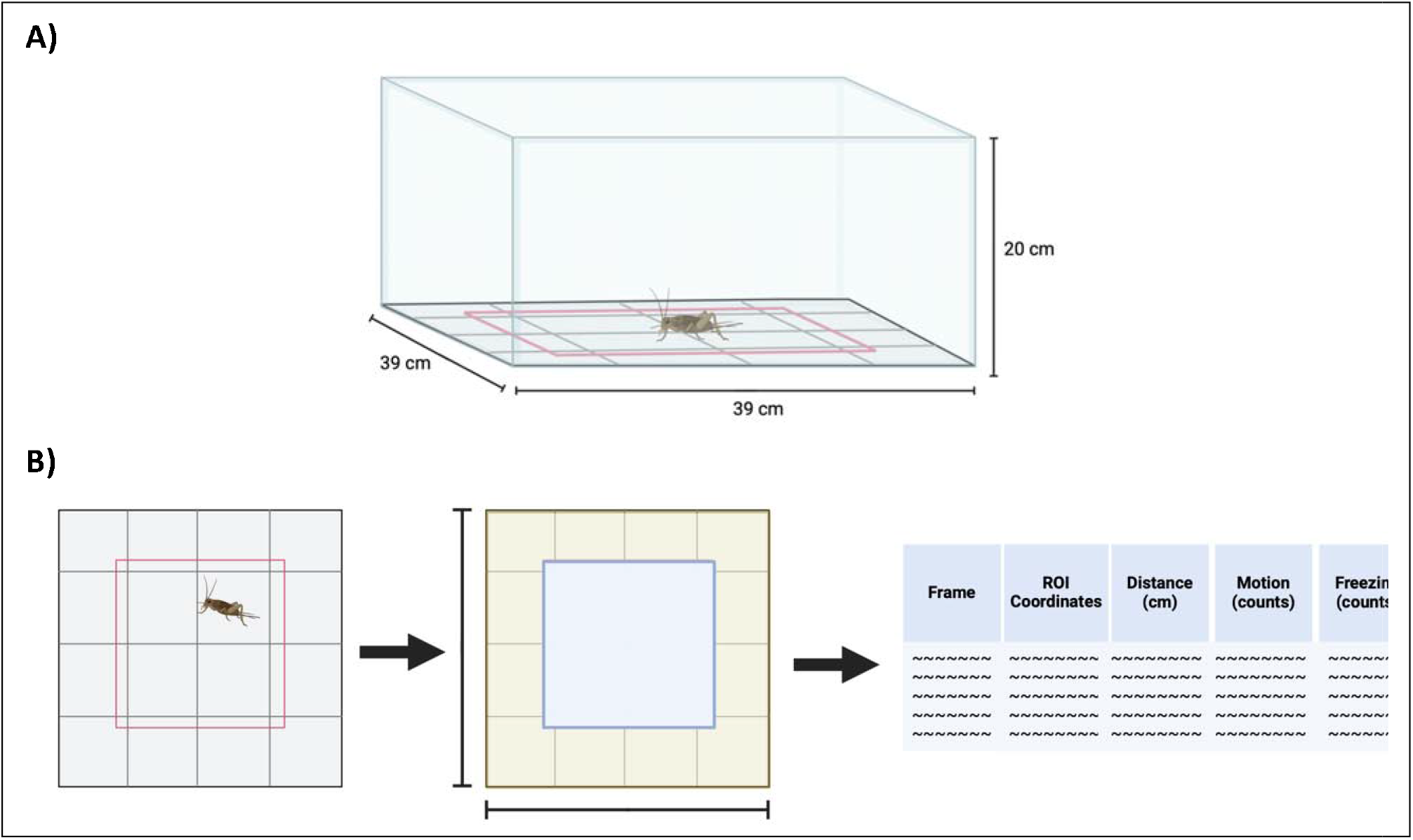
ezTrack open-field test (OFT). **(A)** Schematic diagram of the OFT **(B)** Workflow of ezTrack software for OFT analysis. Figures created using Biorender.com.

Velocity outliers (>99^th^ percentile) were excluded. From processed data, total distance, average speed, freezing events, and time in each zone were derived, with freezing defined as sustained low-velocity motion. Walking and running states were classified using k-means clustering (k = 2) applied to velocity distributions, enabling extraction of state-specific distance, time, and speed. All measures were computed separately for central and peripheral zones.

### Composite frailty index (CFI)

Frailty was quantified using a CFI integrating physical and cognitive behavioral features derived from open-field tracking. Ten parameters were extracted and grouped into physical (total distance, average gait speed, mean walking speed, mean running speed, walking-to-running distance ratio, walking-to-running time ratio) and cognitive domains (central-to-peripheral distance ratio, central-to-peripheral time ratio, total freezing duration, central freezing-to-motion ratio).

To capture deviation from youthful or healthy function, each variable was normalized to a reference population (young adults for aging analyses; age-matched controls for pharmacological studies). Normalized values were discretized into quintiles and assigned ordinal scores (0-4; 0 indicates preserved [youth-like] function, 4 indicates maximal frailty), reducing sensitivity to outliers and ensuring comparable feature weighting.

The CFI was calculated as the sum of all ten feature scores (range: 0-40), with higher values indicating greater integrated frailty and enabling comparison across age groups and treatment conditions.

### Ethics and euthanasia

Experiments were conducted using *Acheta domesticus*, an invertebrate species not subject to IACUC oversight. All procedures adhered to institutional and international guidelines for invertebrate research and followed ARRIVE recommendations where applicable. Crickets were euthanized by controlled CO_2_ exposure followed by decapitation at the cranio-cervical junction, consistent with AVMA guidelines [22].

### Statistical analysis

Continuous variables were presented as mean (standard deviation [SD]). Locomotor states were defined by k-means clustering (k = 2) of velocity distributions, with cluster validity assessed by silhouette scores and visual inspection. Data normality was evaluated using the Shapiro-Wilk test. For sex-stratified group analyses, two-way analysis of variance (ANOVA) was used with Bonferroni or Dunnett’s post hoc correction. Effect sizes were calculated using Cohen’s *d* with Hedges’ *g* correction and 95% confidence intervals (CI), with either young-adult or control as reference. Analyses were performed in GraphPad Prism (v10.0.3; GraphPad Software) and Python (v3.13.1; Python Software Foundation) using SciPy (1.13.1), *scikit-learn*, and matplotlib (v3.9.2). Statistical significance was set at α = 0.05.

## Results

### Velocity-based identification and validation of locomotor states across aging and pharmacological intervention

Unsupervised k-means clustering (k = 2) of locomotor velocity identified two distinct states corresponding to walking and running across all cohorts (Appendix 1, Fig.1). Velocity thresholds were derived from center-of-mass displacement (ezTrack “Location_output” [20]) after exclusion of values above the 99^th^ percentile, yielding consistently separated low- and high-speed distributions.

Across aging, young adults exhibited higher walking (3.35 ± 3.95 cm/s) and running speeds (29.82 ± 2.23 cm/s) than geriatrics (1.97 ± 2.25 cm/s; 21.56 ± 8.31 cm/s), indicating reduced maximal locomotor output with age (Table 1). Pharmacological cohorts showed graded effects, with rapamycin producing the highest velocities (2.93 ± 3.69 cm/s; 30.51 ± 11.64 cm/s) relative to controls (2.73 ± 3.28 cm/s; 28.05 ± 11.22 cm/s), and acarbose and phenylbutyrate showing intermediate reductions (Table 1). Independent validation using motion magnitude (ezTrack “Freezing_output”) confirmed robust cluster separation, with silhouette scores ranging from 0.600 to 0.655 across all groups, indicating strong cluster integrity and minimal overlap between locomotor states (Table 1; Appendix 1, Fig 2.) [23-24].

**Table 1.**
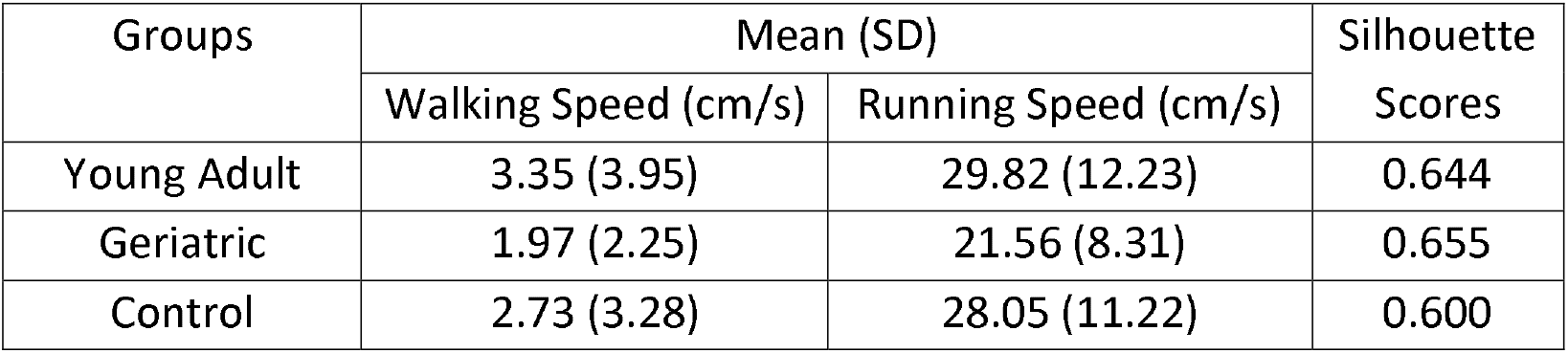
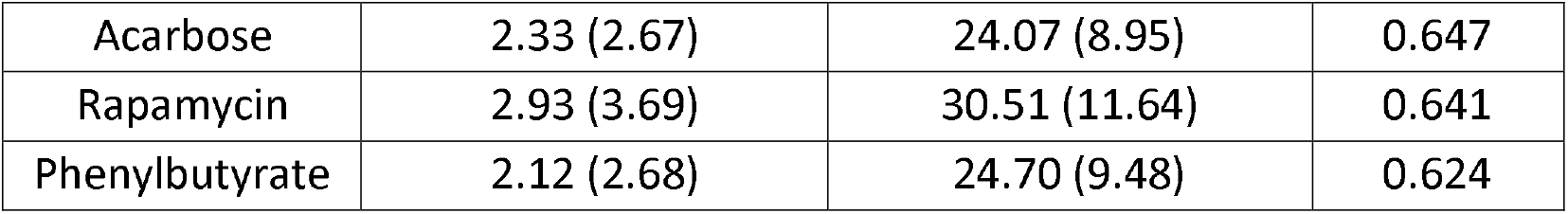
Group-dependent velocity distributions and silhouette-based cluster validation.

**Fig. 2.**
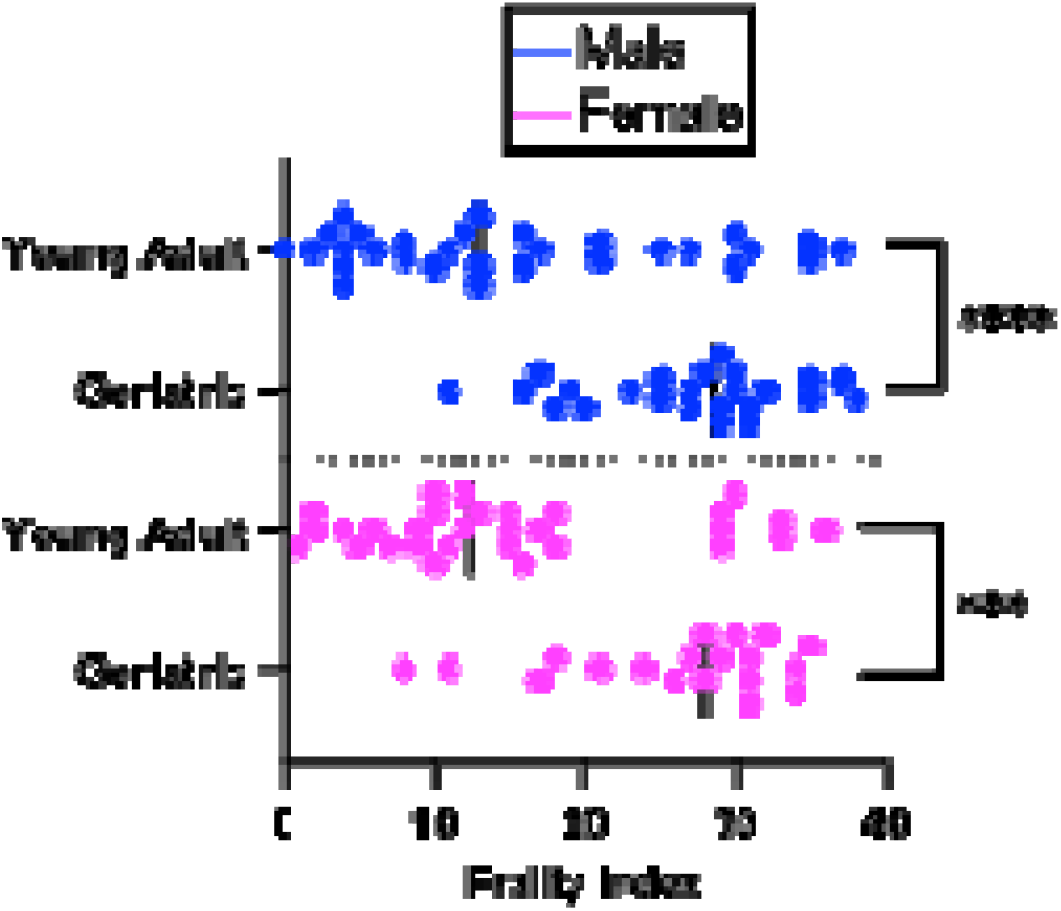
Age-dependent differences in composite frailty. CFI scores stratified by sex across age groups. Each point represents one individual; horizontal lines indicate group medians, ^***^P< 0.001, ^****^P< 0.0001.

### CFI reveals age-dependent functional decline

CFI increased with chronological age, reflecting progressive deterioration of integrated physical and cognitive function (Fig. 2). Young adult females exhibited lower CFI scores (15.20 ± 10.19) compared to geriatric females (26.05 ± 7.74) (*d* = 1.14 [95% CI: 0.53, 1.76], *P* = 0.0003). Males experienced similar differences, with young adults exhibiting lower scores than geriatrics (15.37 ± 11.25 vs. 26.55 ± 7.01; *d* = -1.17 [95% CI: - 1.75 to -0.59], *P* < 0.0001).

CFI trajectories were comparable between sexes, with minimal separation between males and females within each age group (Fig. 2). Variation in CFI was predominantly explained by age (*F* = 34.22, *P* < 0.0001), with negligible contributions from sex (*F* = 0.052, *P* = 0.82) and age-by-sex interaction (*F* = 0.019, *P* = 0.89), indicating that CFI primarily captures age-dependent functional decline independent of sex.

### Pharmacological modulation of frailty reveals treatment-dependent improvement independent of sex

Pharmacological intervention produced treatment-dependent reductions in CFI, with rapamycin yielding the lowest frailty scores across both sexes (Fig. 3). In females, CFI was reduced in rapamycin-treated crickets (11.40 ± 9.73) relative to controls (24.29 ± 9.58) (*d* = -1.31 [95% CI: -2.09, -0.53], *P* = 0.0017). Acarbose (24.00 ± 9.19) or phenylbutyrate-treated males (18.80 ± 11.01) did not differ from controls (*d*’s = -0.54 to -0.03; *P*’s > 0.05). A similar pattern was observed in males, with rapamycin-treated crickets exhibiting lower CFI (9.00 ± 9.63) compared to controls (23.06 ± 10.53) (*d* = -1.33 [95% CI: -2.09, -0.58], *P* = 0.0004), while no differences were observed for acarbose (19.40 ± 11.04), and phenylbutyrate (20.80 ± 6.03) versus controls (*d*’s = -0.34 to -0.23; *P* > 0.05).

**Fig. 3.**
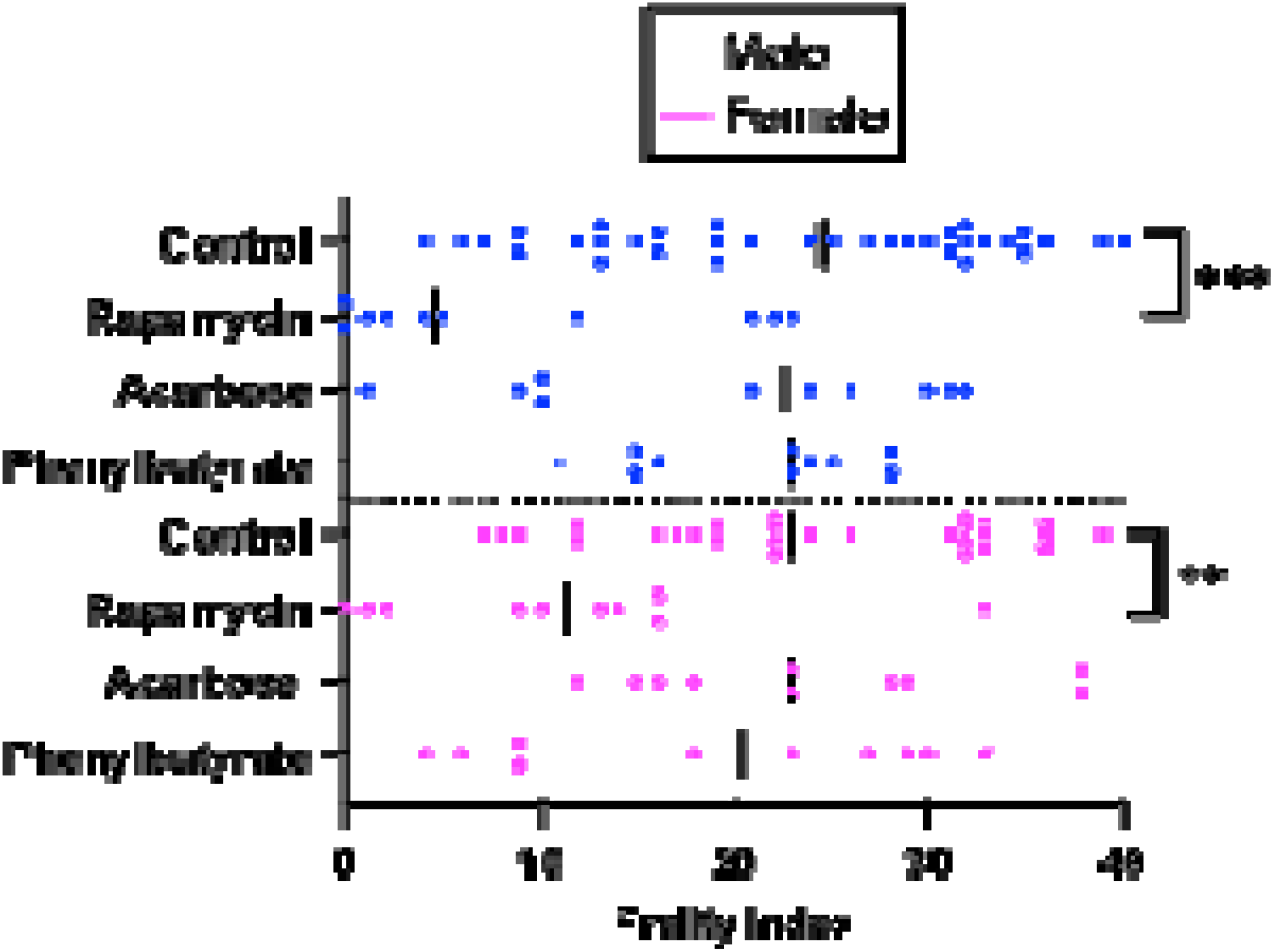
Pharmacological attenuation of late-life frailty. Composite Frailty Index (CFI) scores stratified by sex across treatment groups. Each point represents one individual; horizontal lines indicate group medians, ^**^P< 0.01, ^***^P< 0.001.

Frailty reduction was comparable between males and females, with minimal divergence in treatment response (Fig. 3). Variation in CFI was primarily driven by treatment group (*F* = 9.512, *P* < 0.0001), with negligible contributions from sex (*F* = 0.599, *P* = 0.441) and treatment-by-sex interaction (*F* = 0.392, *P* = 0.759), indicating that pharmacologic modulation of functional aging is largely sex independent.

## Discussion

A central challenge in geroscience is that experimental endpoints often remain misaligned with the clinical reality of aging. While lifespan is a powerful biological measure, it does not capture the progressive loss of physical, cognitive, and physiological resilience to stress that defines frailty and determines late-life outcomes [25]. Here, we addressed this gap by introducing the first integrative frailty index in an invertebrate, establishing the house cricket (*Acheta domesticus*) as a scalable model for quantifying multidimensional functional decline.

The CFI captures coordinated deterioration across locomotion, exploration, and freezing behavior, generating a continuous phenotype that increases with age and responds to intervention. Using this framework, we showed that mid-life rapamycin treatment attenuates late-life frailty, indicating preservation of integrated physical and cognitive function. In contrast, acarbose and phenylbutyrate produced inconclusive effects under the conditions tested. These findings highlight the ability of this system to prioritize candidate geroprotectors based on functional outcomes rather than lifespan alone, despite all three compounds having established clinical use (rapamycin as an immunosuppressant [26], acarbose for glycemic control [27], and phenylbutyrate for metabolic disorders [28]).

Importantly, the CFI is designed as a first-pass screening tool within a hierarchical pipeline rather than a comprehensive assessment of aging. Compounds that fail to improve frailty in this initial screen will be retested across multiple doses and larger cohorts, while those that demonstrate consistent benefit will advance to deeper phenotyping, including geropathology, expanded cognitive assays [17], and pharmacokinetic profiling. This staged approach addresses a major bottleneck in geroscience by enabling rapid prioritization of candidate interventions for downstream validation [29].

Beyond pharmacological screening, this platform enables integration of functional decline with tissue-level pathology in a high-throughput system. The extensibility of the CFI framework allows systemic evaluation of a broad range of repurposed gerotherapeutics, including SGLT2 inhibitors, metformin, GLP-1 receptor agonists and bisphosphonates [30]. This creates a scalable, continuously adaptable discovery platform capable of aligning experimental aging biology with clinically meaningful outcomes.

Several limitations warrant consideration. The current CFI emphasizes gross locomotor and behavioral features and does not capture fine-scale motor dynamics such as stride variability or inter-limb coordination, which may represent earlier indicators of decline [31]. In addition, sample sizes were constrained by feasibility, limiting sensitivity to smaller effect sizes. Future studies incorporating high-resolution behavioral tracking [32-33] and larger, multi-site cohorts will further enhance the precision and translational relevance of this system.

## Conclusion

This work establishes the house cricket (*Acheta domesticus*) as a function-centered, high-throughput model of aging with frailty as a primary endpoint. By positioning the CFI within a layered geroscience pipeline, this platform enables rapid identification of interventions that preserve not only lifespan, but functional healthspan.

## Supporting information

Biorender Publication License

Biorender Publication License

## Acknowledgments

This work was supported by the National Institute on Aging of the National Institutes of Health under award number R01 AG067193. The funder had no role in study design, data collection and analysis, decision to publish, or preparation of the manuscript.

## Availability of Data and Materials

The datasets generated during the current study are available from the corresponding author on reasonable request.

## References

1. Clegg A, Young J, Iliffe S, Rikkert MO, Rockwood K. Frailty in elderly people. Lancet. 2013;381(9868):752–762. doi:10.1016/S0140-6736(12)62167-9

2. Bergman H, Ferrucci L, Guralnik J, et al. Frailty: an emerging research and clinical paradigm--issues and controversies. J Gerontol A Biol Sci Med Sci. 2007;62(7):731–737. doi:10.1093/gerona/62.7.731

3. Gonçalves RSDSA, Maciel ÁCC, Rolland Y, Vellas B, de Souto Barreto P. Frailty biomarkers under the perspective of geroscience: A narrative review. Ageing Res Rev. 2022;81:101737. doi:10.1016/j.arr.2022.101737

4. Kulkarni AS, Aleksic S, Berger DM, Sierra F, Kuchel GA, Barzilai N. Geroscience-guided repurposing of FDA-approved drugs to target aging: A proposed process and prioritization. Aging Cell. 2022;21(4):e13596. doi:10.1111/acel.13596

5. Palliyaguru DL, Moats JM, Di Germanio C, Bernier M, de Cabo R. Frailty index as a biomarker of lifespan and healthspan: Focus on pharmacological interventions. Mech Ageing Dev. 2019;180:42–48. doi:10.1016/j.mad.2019.03.005

6. Mitchell SJ, Bernier M, Aon MA, et al. Animal models of aging research: implications for human healthspan. Nat Aging. 2025;5(1):149–162. doi:10.1038/s43587-025-00149-y

7. Andersen, Monica L, and Lucile M F Winter. “Animal models in biological and biomedical research - experimental and ethical concerns.” Anais da Academia Brasileira de Ciencias vol. 91, suppl 1 (2019): e20170238. doi:10.1590/0001-3765201720170238

8. Partridge L, Tower J (2007) Yeast a feast: the fruit fly Drosophila as a model organism for research into aging. In: Guarente L, Partridge L, Wallace DC (eds). Mol Biol Aging. Cold Spring Harbor Monograph Series 51:267–308

9. Lyn JC, Naikkhwah W, Aksenov V, & Rollo CD. Influence of two methods of dietary restriction on life history features and aging of the cricket Acheta domesticus. Age (Dordr), 2011, 33(4): 509–522.

10. Litke, Rachel et al. “Caenorhabditis elegans, un modèle d’étude du vieillissement” [Caenorhabditis elegans as a model organism for aging: relevance, limitations and future]. Medecine sciences : M/S vol. 34, 6-7 (2018): 571–579. doi:10.1051/medsci/20183406017

11. Liao GY, Dai S, Bae E, et al. Morphological features of the domestic house cricket (Acheta domesticus) for translational aging studies. Geroscience. Published online June 3, 2025. doi:10.1007/s11357-025-01711-9

12. Liao GY, Rosenfeld M, Wezeman J, Ladiges W. The house cricket is an unrecognized but potentially powerful model for aging intervention studies. Aging Pathobiol Ther. 2024;6(1):39–41. doi:10.31491/apt.2024.03.135

13. Pitt JN, Kaeberlein M. Why is aging conserved and what can we do about it? [published correction appears in PLoS Biol. 2015 May 15;13(5):e1002176. doi: 10.1371/journal.pbio.1002176.]. PLoS Biol. 2015;13(4):e1002131. Published 2015 Apr 29. doi:10.1371/journal.pbio.1002131

14. Ghosal K, Gupta M, Killian KA. Agonistic behavior enhances adult neurogenesis in male Acheta domesticus crickets. J Exp Biol. 2009;212(Pt 13):2045–2056. doi:10.1242/jeb.026682

15. Dobson GP, Letson HL, Biros E, Morris J. Specific pathogen-free (SPF) animal status as a variable in biomedical research: Have we come full circle?. EBioMedicine. 2019;41:42–43. doi:10.1016/j.ebiom.2019.02.038

16. Bundgaard CJ, Kalliokoski O, Abelson KS, Hau J. Acclimatization of mice to different cage types and social groupings with respect to fecal secretion of IgA and corticosterone metabolites. In Vivo. 2012;26(6):883–888.

17. Liao GY, Klug J, Ladiges WC. Age-related cognitive decline in house crickets reveals conserved patterns of sensory and learning deficits across the lifespan. bioRxiv. Posted August 14, 2025. doi:10.1101/2025.08.14.670304.

18. Liao GY, Klug J, Singh S, Bae E, Dai S, Ladiges W. The gerotherapeutic drugs rapamycin, acarbose, and phenylbutyrate extend lifespan and enhance healthy aging in house crickets. Preprint. Res Sq. 2026;rs.3.rs-7466146. Published 2026 Jan 5. doi:10.21203/rs.3.rs-7466146/v1

19. Jiang Z, Wang J, Imai D, et al. Short term treatment with a cocktail of rapamycin, acarbose and phenylbutyrate delays aging phenotypes in mice. Sci Rep. 2022;12(1):7300. Published 2022 May 4. doi:10.1038/s41598-022-11229-1

20. Pennington, Z. T., Dong, Z., Feng, Y., Vetere, L. M., Page-Harley, L., Shuman, T., & Cai, D. J. (2019). ezTrack: An open-source video analysis pipeline for the investigation of animal behavior. Scientific Reports, 9(1), 19979.

21. McElroy DL, Roebuck AJ, Onofrychuk TJ, Sandini TM, Greba Q, Howland JG. Implementation of ezTrack open-source pipeline for quantifying rat locomotor behavior: Comparison to commercially available software. Neurosci Lett. 2020;723:134839. doi:10.1016/j.neulet.2020.134839

22. Leary S. Guidelines for the euthanasia of animals. American Veterinary Medical Association. 2020. Accessed February 25, 2025. https://www.avma.org/resources-tools/avma-policies/avma-guidelines-euthanasia-animals.

23. Rousseeuw PJ. Silhouettes: A graphical aid to the interpretation and validation of cluster analysis. Journal of Computational and Applied Mathematics. 1987;20:53–65. doi:10.1016/0377-0427(87)90125-7

24. Dalmaijer ES, Nord CL, Astle DE. Statistical power for cluster analysis. BMC Bioinformatics. 2022;23(1):205. Published 2022 May 31. doi:10.1186/s12859-022-04675-1

25. de Breij S, Rijnhart JJM, Schuster NA, Rietman ML, Peters MJL, Hoogendijk EO. Explaining the association between frailty and mortality in older adults: The mediating role of lifestyle, social, psychological, cognitive, and physical factors. Prev Med Rep. 2021;24:101589. Published 2021 Oct 7. doi:10.1016/j.pmedr.2021.101589

26. Roark KM, Iffland PH 2nd. Rapamycin for longevity: the pros, the cons, and future perspectives. Front Aging. 2025;6:1628187. Published 2025 Jun 20. doi:10.3389/fragi.2025.1628187

27. McIver LA, Preuss CV, Tripp J. Acarbose. [Updated 2024 Feb 12]. In: StatPearls [Internet]. Treasure Island (FL): StatPearls Publishing; 2025 Jan-. Available from: https://www.ncbi.nlm.nih.gov/books/NBK493214/

28. LiverTox: Clinical and Research Information on Drug-Induced Liver Injury [Internet]. Bethesda (MD): National Institute of Diabetes and Digestive and Kidney Diseases; 2012-. Phenylbutyrate, Sodium Benzoate. [Updated 2016 Oct 10]. Available from: https://www.ncbi.nlm.nih.gov/books/NBK547972/

29. Moskalev A, Chernyagina E, de Magalhães JP, et al. Geroprotectors.org: a new, structured and curated database of current therapeutic interventions in aging and age-related disease. Aging (Albany NY). 2015;7(9):616–628. doi:10.18632/aging.100799

30. Leone, M., & Barzilai, N. (2024). An Updated Prioritization of Geroscience-Guided FDA-Approved Drugs Repurposed to Target Aging. Medical Research Archives, 12(2). doi:10.18103/mra.v12i2.5138

31. Plotnik M, Giladi N, Hausdorff JM. A new measure for quantifying the bilateral coordination of human gait: effects of aging and Parkinson’s disease. Exp Brain Res. 2007;181(4):561–570. doi:10.1007/s00221-007-0955-7

32. Barreto L, Shon A, Knox D, Song H, Park H, Kim J. Motorized Treadmill and Optical Recording System for Gait Analysis of Grasshoppers. Sensors. 2021; 21(17):5953. 10.3390/s21175953

33. Mock JT, Knight SG, Vann PH, et al. Gait Analyses in Mice: Effects of Age and Glutathione Deficiency. Aging Dis. 2018;9(4):634–646. Published 2018 Aug 1. doi:10.14336/AD.2017.0925

