## Appendix 1 for "A composite frailty index enables quantification of functional aging and identification of gerotherapeutic drugs in the house cricket"

**Appendix 1. Velocity distributions, silhouette plots, and cluster visualizations.**

**A)	B)**

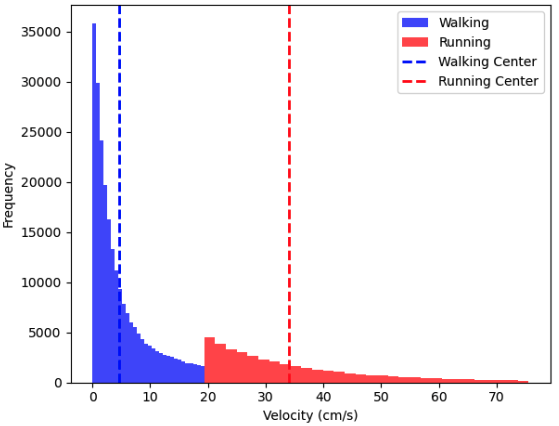

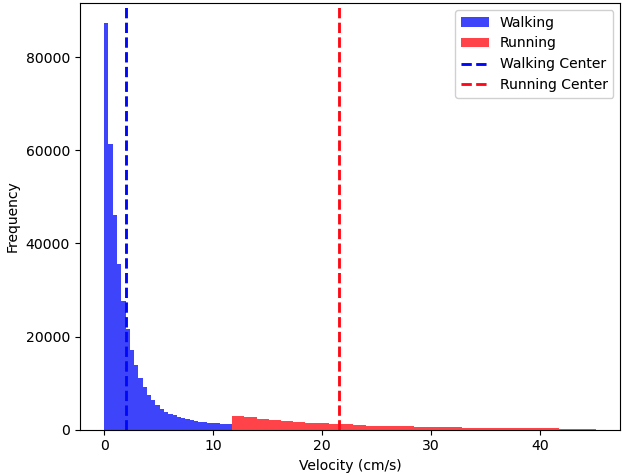

**C)	D)**

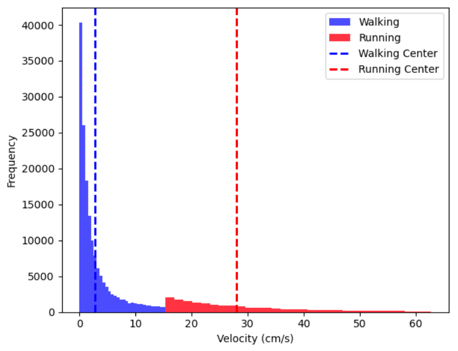

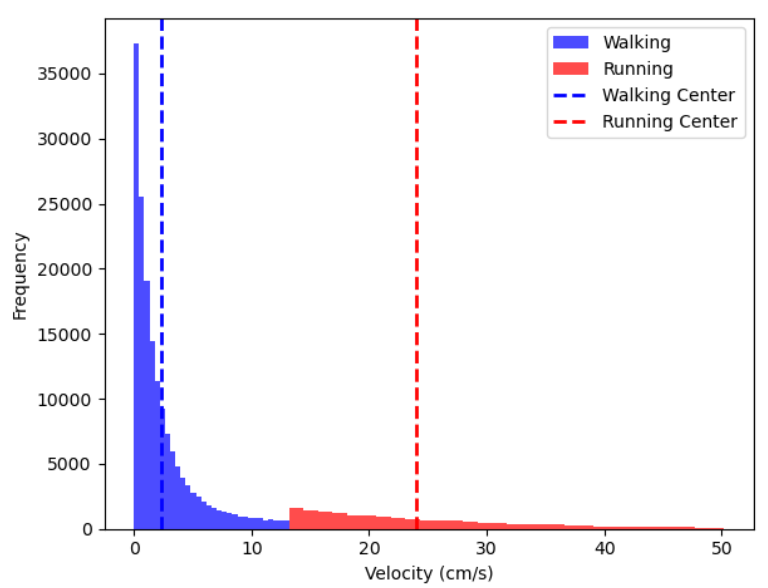

**E)	F)**

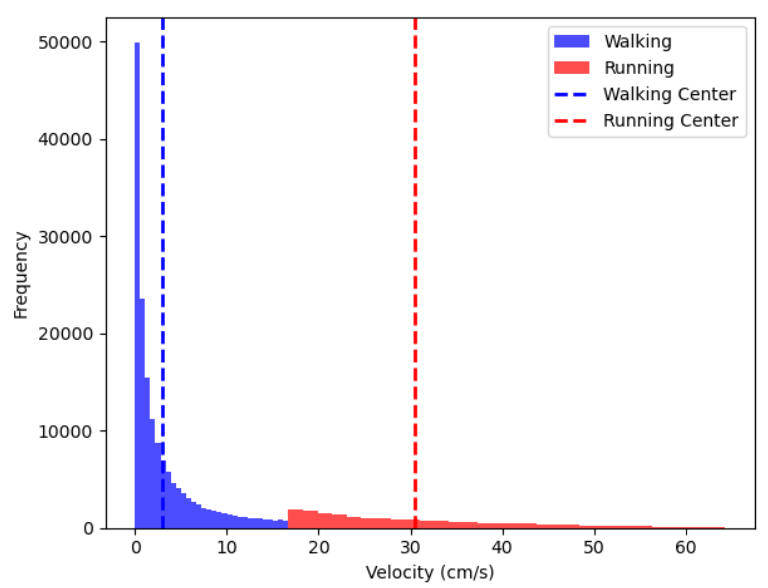

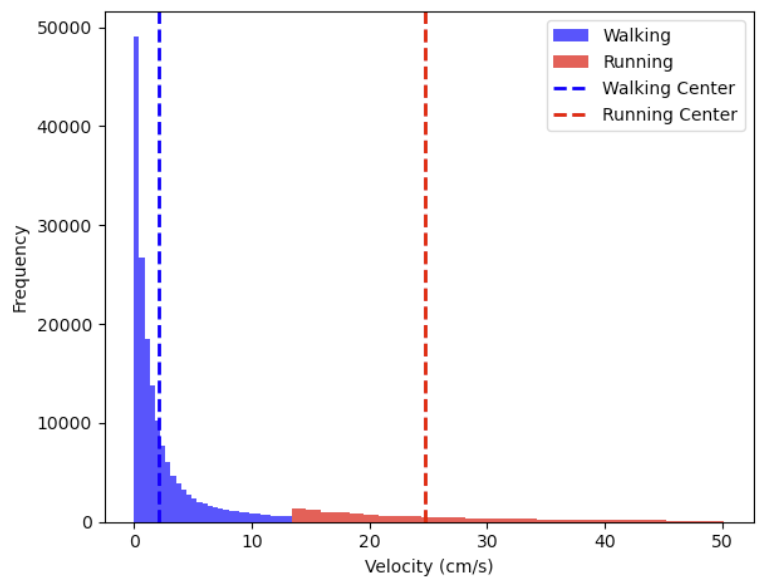

**Figure 1. Velocity-defined locomotor states across aging and treatment groups. (A-B)** Probability density plots of center-of-mass velocity (cm/s) for **(A)** young-adult and **(B)** geriatric crickets. **(C-F)** Velocity distributions for **(C)** control, **(D)** acarbose, **(E)** rapamycin, and **(F)** phenylbutyrate cohorts. Vertical dashed lines indicate k-means cluster centroids separating walking and running states.

**A)**

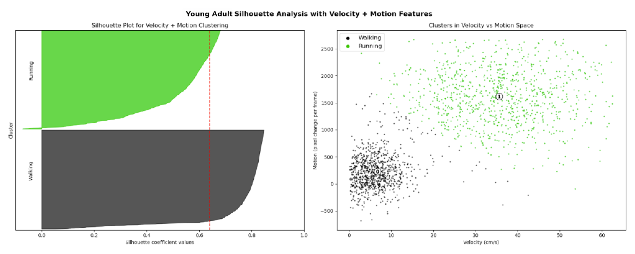
**B)**

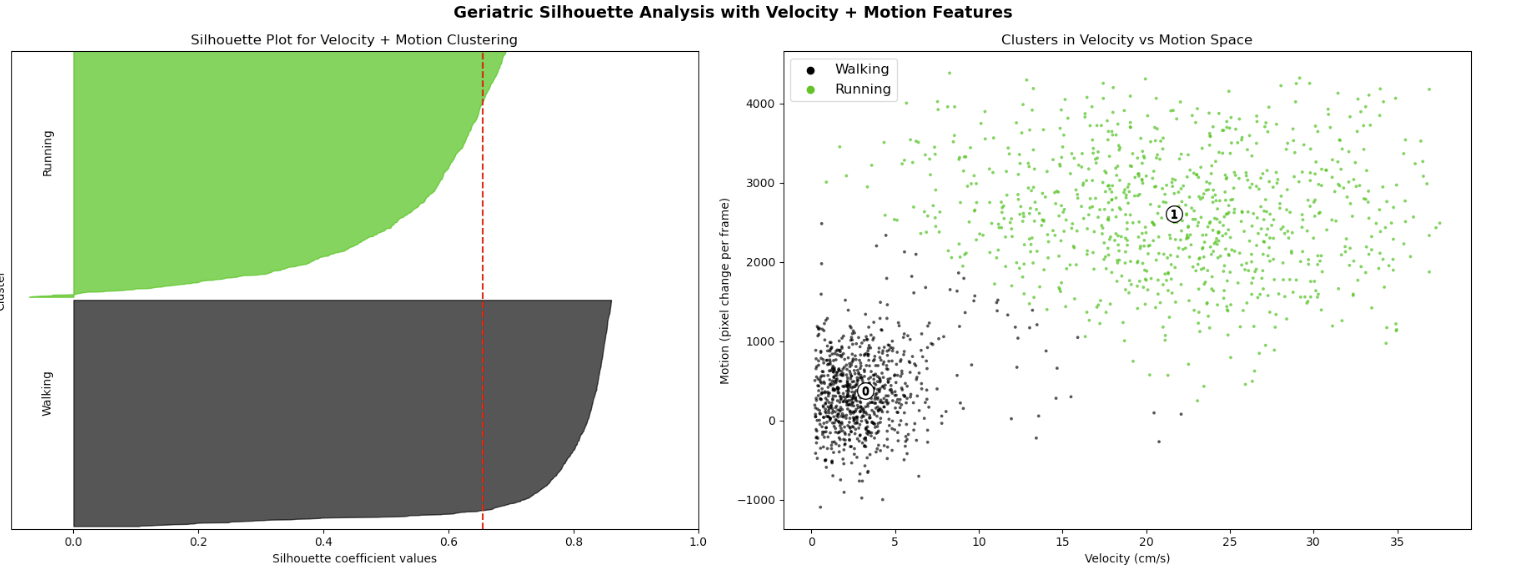

**C)**

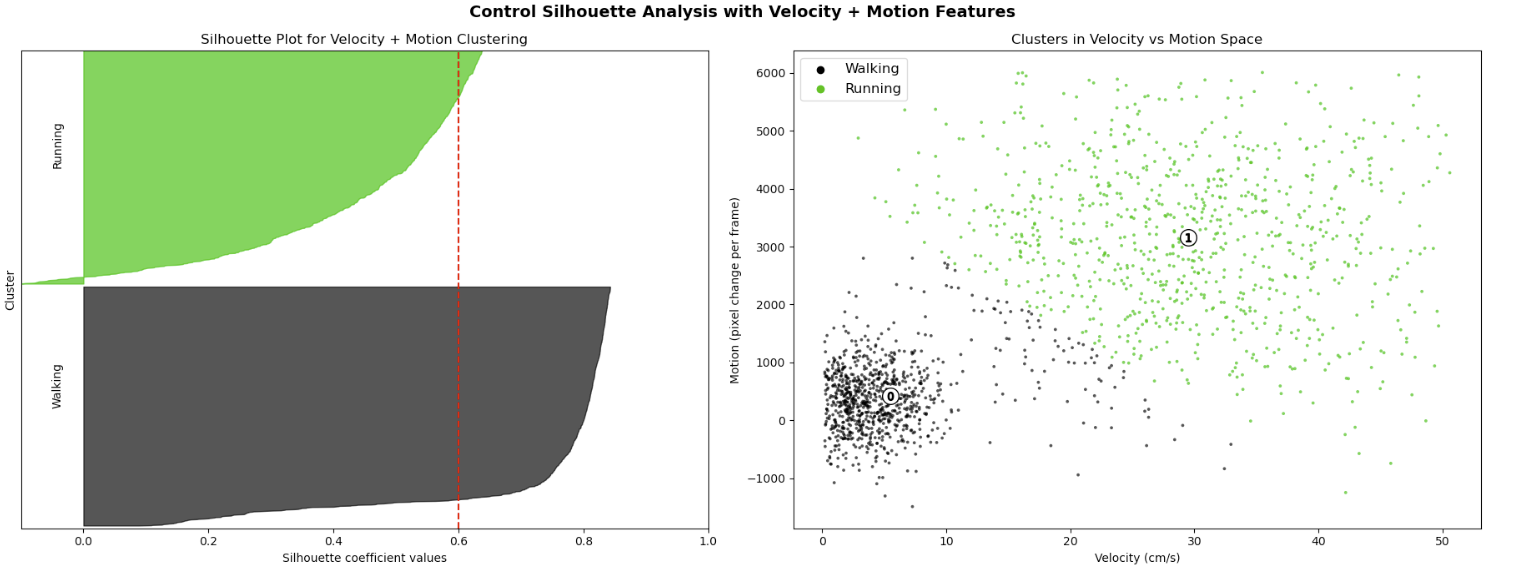
**D)**

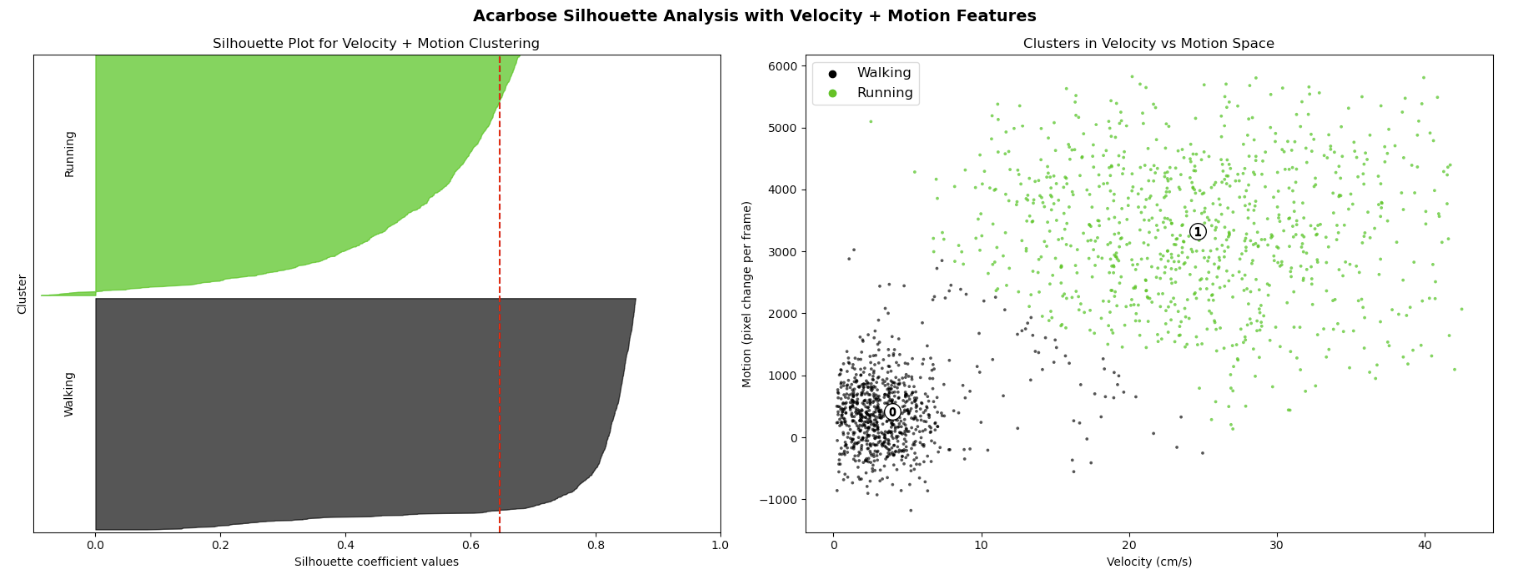

**E)**
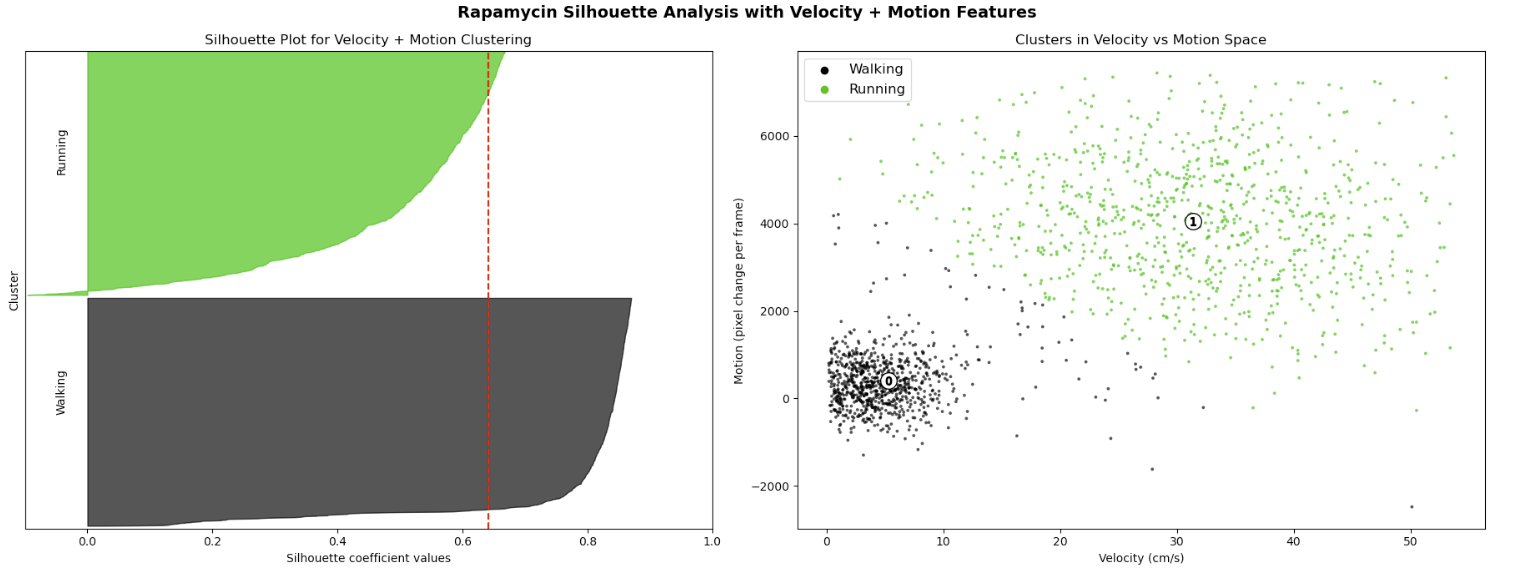

**F)**
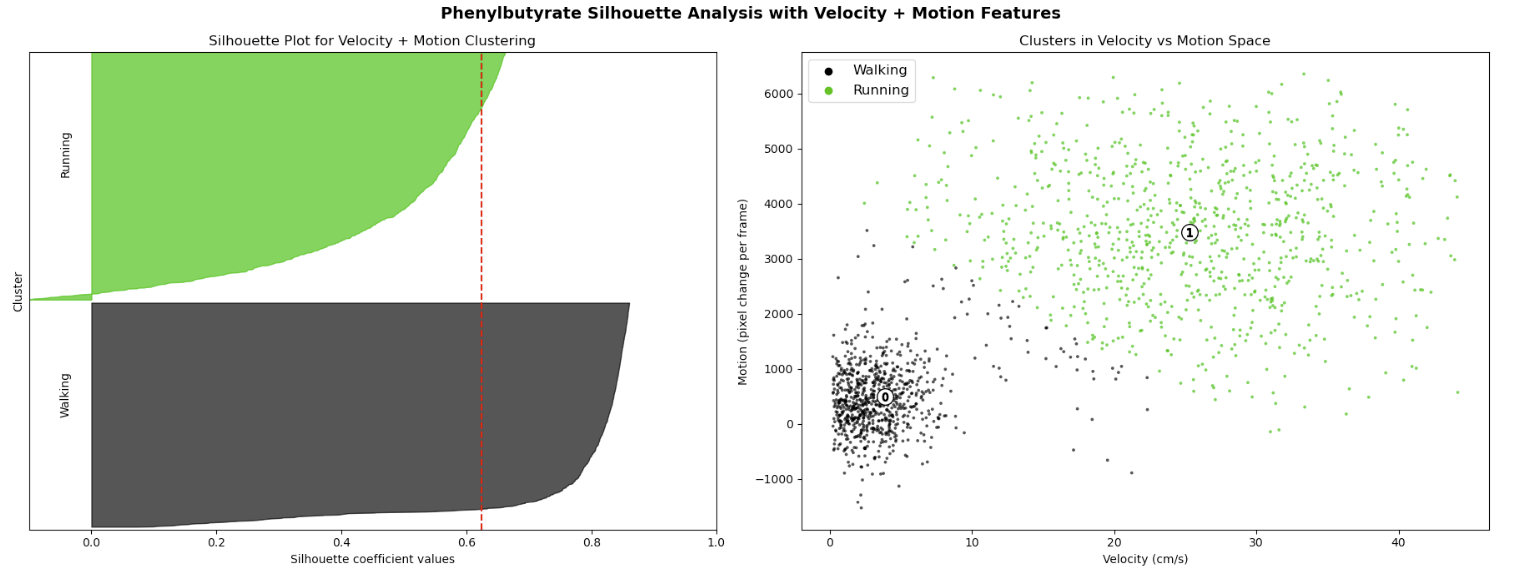

**Figure 2. Silhouette-based validation across aging and treatment groups. (A-F)** Two-dimensional k-means clustering of velocity and motion magnitude metrics for **(A)** young-adult, **(B)** geriatric, **(C)** control, **(D)** acarbose, **(E)** rapamycin, and **(F)** phenylbutyrate cohorts. Left: Silhouette plots showing cluster compactness and separation; Right: scatter plots showing separation of walking and running datapoints. In all age and treatment conditions, mean silhouette scores (red vertical lines) exceeded 0.5.
