## Supplementary material for "A composite frailty index enables quantification of functional aging and identification of gerotherapeutic drugs in the house cricket": Biorender Publication License

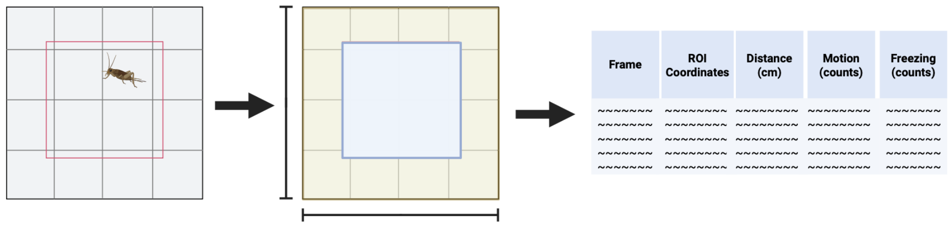


*For any questions regarding this document, or other questions about publishing with BioRender, please refer to our BioRender Publication Guide, or contact BioRender Support at.*
